# Retrospective Study on the Health Problems of Falcons in Al Ain, United Arab Emirates

**DOI:** 10.1101/2021.02.11.430749

**Authors:** Maryam Abdullah Al Hemeiri, Abraham Arias de la Torre, Khaja Mohteshamuddin, Berhanu Adenew Degafa, Gobena Ameni

**Affiliations:** Department of Veterinary Medicine, College of Food and Agriculture, United Arab Emirates University, PO Box 15551, Al Ain, UAE; City Vet Clinic, Al Bawadi Market Enterprise, Al Ain City, Abu Dhabi, UAE; Department of Integrative Agriculture, United Arab Emirates University, PO Box 15551, Al Ain, UAE; Aklilu Lemma Institute of Pathobiology, Addis Ababa University, PO box 1176, Addis Ababa, Ethiopia

**Keywords:** Aspergillosis, Bacterial enteritis. Falcon health, Ingluvitis, United Arab Emirates

## Abstract

**Background:** Falcons are important animals in sociocultural events of the society of the United Arab Emirates (UAE). Like any other birds, falcons can be affected by different health problems. This study was conducted to investigate the major health problems of falcons presented to the City Vet Clinic in Al Ain during 2019.

**Methods:** Data were extracted from the records of 906 falcons presented to City Vet Clinic in 2019. Data extraction was made on the diagnostic methods used, diagnosed health problems and the type of treatment/veterinary services given to falcons. Analysis was done using descriptive statistics.

**Results:** The overall incidence rate of health problems detected in falcons presented to City Vet Clinic in 2019 was 26.5% (95% confidence interval, CI, 23.6-29.5%). The most incident health problems were ingluvitis (inflammation of crop), aspergillosis and bacterial enteritis with incidence rates of 8.1% (95%CI: 6.4–10.0%), 5.8% (95%CI: 4.4-7.6%) and 2.4% (95%CI: 1.5-3.7%), respectively. The relationship between the number of cases of falcon and months was polynomial with a regression (R^2^) of 42% indicating that only 42% of the variation in the number cases could be explained by monthly variation. The three main medical treatments given for the diagnosed health problems included antibiotics, anti-fungal and anti-parasitic with frequencies of 46.3% (95%CI: 39.8-52.8%), 21.7% (95%CI: 16.6-27.4%) and 12.0% (95%CI: 8.2-16.9%), respectively.

**Conclusion:** The major health problems of falcons were ingluvitis, aspergillosis and bacterial enteritis. Infections that occur in falcons can also be transmitted to owners. Therefore, regular check-up and control of diseases of falcons is recommended.

## Background

Falcons are small to medium size, strong and rigid birds which are characterized by their swift, graceful and predatory skill [1,2]. Taxonomically, falcons belong to the Phylum Chordata, Family Falconinae, Subfamily Falconidae, Order Falconiformes and the Genus Falco [3]. There are over 35 species in the Genus *Falco* and they are distributed worldwide [2]. The most common species of the Genus Falco, which are found in the United Arab Emirates (UAE) are *Falco cherrug* (Saker), *Falco rusticolus* (Gyrfalcon), *Falco peregrinus* (Peregrine), hybrid of Saker-Gyrfalcon, and hybrid of Peregrine-Gyrfalcon [4].

Falconry is also termed as hawking is a type of sport, which employs primarily falcons and to certain extent, eagles or buzzards in hunting game [5]. The term falconry is directly linked to falcons as they are the primary birds involved in this sport and for which reason they are termed as true hawks [5]. In the UAE and other Middle East countries, the role of falcons has not only limited to sports and entertainment but also, they have special places in the tradition of the societies [6]. The special place given to the falcons by the Emirati people and other people of the Middle East originated from the historical relationship between the Bedouins and the falcons [7,8]. Bedouins are Arabic-speaking nomadic peoples of the Middle Eastern deserts, especially of North Africa, the Arabian Peninsula, Egypt, Israel, Iraq, Syria, and Jordan in the past [9]. Bedouins were living in the desert under harsh environment and thus, they used to trap wild falcons passing over the sky the of UAE and the Arabian Peninsula on the migratory route from Europe to Africa and train them for two weeks and used for hunting migratory birds [8]. The preys that were hunted by falcons and cooked as meals for Bedouins families [8]. Therefore, falconry was a matter of necessity for Bedouins to survive under harsh conditions in the desert, which lead the Emirati people to have a different view for falcons in which they do not regard falcons as only sport animals like the European but also consider them as the integral part of their families [4]. The Emirati people do not value falcons in terms of money; but rather they value them in the same way they value their sons and daughters, and as a result, falcons are living in a well-ventilated and furnished rooms, have their own places in their owners’ cars and even in offices [6]. Falconry is a traditional sport, attracts the leaders and the ordinary people. The new generations of the Emirati people have inherited falconry from their ancestors and hence falconry is considered as the most common tradition of the UAE people [4]. Reports show that training of falcons for the legendary sport costs a huge amount of money and according to Jacobs [10], in the UAE, falconers train their falcons, which can cost up to $60,000 pet a bird, to race at hundreds of miles an hour in the President’s Cup, a national competition in which the fastest falcons can win up to $7 million in prizes.

Falcons like any other birds can be affected by different health problems. Falcons can be infected by bacterial infections such as salmonellosis and parasitic diseases such as trichomoniasis and coccidiosis [4,11]. Besides, fungal infections such as aspergillosis and viral diseases including avian pox can affect falcons and result in morbidities and mortalities [12]. Moreover, falcons can be infested by external parasites including mites and lice [11].

Therefore, the prevailing socio-cultural situations of the UAE require adequate maintenance of the health of the falcons for two reasons; firstly, falcons are precious animals for the Emirate people and hence, they should be kept healthy and safe, and secondly, because of the strong physical contact between falcons and their owners, there could be a chance of transmission of zoonotic diseases from falcons to their owners, which necessities strict maintenance of the health of falcons. To this effect, generation of epidemiological data on the diseases that affect falcons in UAE is important as it provides evidence to the control authorities and to the owners for appropriate actions. The present study was conducted to identify major health problems of falcons that were presented to City Vet Clinic in Al Ain, UAE in 2019.

## Materials and methods

### Study setting

The study was conducted at City Vet Clinic, which is located in Al Ain, the United Arab Emirate. The Clinic is situated in the Al Bawadi District of the Al Ain City and is one the Veterinary Clinics treating falcon’s health problem. It is the most popular and famous clinic with a large number of clientele base with regard to falcons. It also deals with the health problems of other exotic species and small animals. The Clinic is well equipped with advanced diagnostic tools. Furthermore, the Veterinary Clinicians working in the City Vet Clinic are highly skilled and well experienced in Falcon Medicine.

### Study design and data collection

The design of this study was a cross-sectional in which retrospective longitudinally recorded data of health problems of falcons were extracted from City Vet Clinic records and analyzed. The City Vet Clinic was first approached and the plan of the study was explained to the Senior Vet Clinician and owner of the Clinic. The Head of the Clinic welcomed the plan and permitted the investigator to use the computerized health data of falcons. The health data of 906 falcon were extracted from the Microsoft Word file of the Clinic and then the data were entered into Microsoft Excel. The extracted data included reasons for which the owners bring their falcons to the clinic, the diagnostic method used by the clinic, the detected falcon health problems, and the treatment given to falcons.

### Data analysis

The data were collected from the City Vet Clinic’s website and re-entered into the Microsoft Excel and then analyzed using descriptive statistics. The overall incidence rate was calculated by dividing the sum of number of falcons affected by the different health problems during 2019 by the total number of falcons visited the City Vet Clinic for different purposes during the same year. The same method of calculation was used to calculate the incidence of each health problem that affected falcons in 2019. Proportions were used to show the frequencies of the different reasons for bringing falcons to the Clinic, the different methods used for the diagnosis and the different types of treatments used. Different Tables were used presenting the summaries of the different aspects of the data. The relationship between the number of cases and months of the Year was assessed using regression analysis and 95% confidence interval was calculated for the incidence rates and frequencies for the evaluation of statistical significance. Incidence rates or proportions having no overlapping confidence intervals were considered as statistically different.

## Results

### Purpose of visit of falcon to the City Vet Clinic

A total of 906 falcons visited the City Vet Clinic for various reasons (Table 1). The majority of falcons (68.9%) were brought to the Clinic for the check-up of their health conditions. Following check-up for health conditions, 11.6% and 8.6% of falcons were brought to the City Vet Clinic for tail mounting and feather fixing, respectively.

**Table 1.**
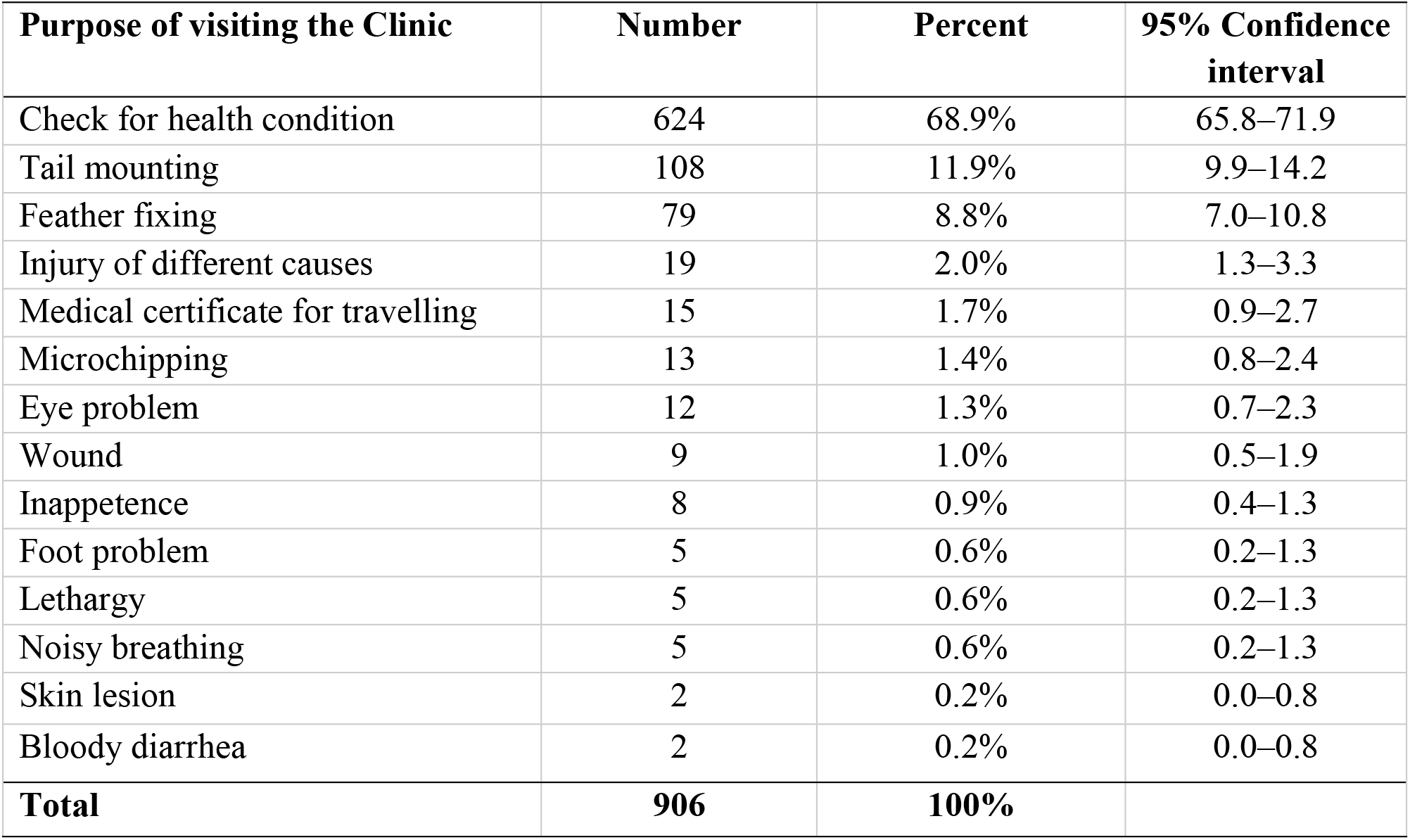
Purposes of owners for taking their falcons to City Vet Clinic in Al Ain City.

| **Purpose of visiting the Clinic** | **Number** | **Percent** | **95% Confidence interval** |
| --- | --- | --- | --- |
| Check for health condition | 624 | 68.9% | 65.8–71.9 |
| Tail mounting | 108 | 11.9% | 9.9–14.2 |
| Feather fixing | 79 | 8.8% | 7.0–10.8 |
| Injury of different causes | 19 | 2.0% | 1.3–3.3 |
| Medical certificate for travelling | 15 | 1.7% | 0.9–2.7 |
| Microchipping | 13 | 1.4% | 0.8–2.4 |
| Eye problem | 12 | 1.3% | 0.7–2.3 |
| Wound | 9 | 1.0% | 0.5–1.9 |
| Inappetence | 8 | 0.9% | 0.4–1.3 |
| Foot problem | 5 | 0.6% | 0.2–1.3 |
| Lethargy | 5 | 0.6% | 0.2–1.3 |
| Noisy breathing | 5 | 0.6% | 0.2–1.3 |
| Skin lesion | 2 | 0.2% | 0.0–0.8 |
| Bloody diarrhea | 2 | 0.2% | 0.0–0.8 |
| **Total** | **906** | **100%** |  |

### Diagnostic methods at the City Vet Clinic for diagnosing falcons

City Vet Clinic used different diagnostic methods for the diagnosis of the health problems of falcons. General clinical observation and imaging of the internal organs were used. In most of the cases, combinations of different diagnostic tools were applied to identify the health problems of falcons. As presented in Table 2, a combination fecal examination and crop endoscopy was the most frequently (32.5%) used diagnostic method in the diagnosis falcon health problem at the City Vet Clinic. Next to the combination of the fecal examination and crop endoscopy, general clinical examination and a combination of three tests (fecal examination, endoscopy and crop endoscopy) were most frequently used and as such 14.1% and 11.9% of falcons were examined by these methods, respectively (Table 2). Fecal examination alone, endoscopy alone, crop endoscopy alone, and X-ray alone were used in 11.8%, 9.9%, 5.4% and 3.1% of the falcons examined at the City Vet Clinic in 2019, respectively.

**Table 2.** Diagnostic methods used for the detection health problems of falcons at the City Vet Clinic, Al Ain City.

| **Diagnostic methods** | **Number examined** | **Percent examined** | **95% Confidence interval** |
| --- | --- | --- | --- |
| Joint fecal and crop endoscopic examinations | 294 | 32.5% | 29.4–35.6 |
| General clinical examination | 126 | 14.1% | 11.7–16.3 |
| Joint fecal, endoscopic and crop endoscopic examinations | 108 | 11.9% | 9.9–14.2 |
| Fecal examination alone | 107 | 11.8% | 9.8–14.1 |
| Endoscopy alone | 90 | 9.9% | 8.1–12.1 |
| Joint endoscopic and crop endoscopic examinations | 51 | 5.6% | 4.2–7.3 |
| Crop endoscopy alone | 49 | 5.4% | 4.0–7.9 |
| X-ray alone | 28 | 3.1% | 2.1–4.4 |
| Joint fecal and endoscopic examinations | 11 | 1.2% | 0.6–2.2 |
| Joint fecal, crop endoscopic and X-ray examinations | 9 | 1.0% | 0.5–1.9 |
| Fluorescent eye stain test | 8 | 0.9% | 0.4–1.7 |
| Joint endoscopic and X-ray examinations | 5 | 0.6% | 0.2–1.3 |
| Joint crop endoscopic, biochemistry and blood tests | 3 | 0.3% | 0.1–1.0 |
| Joint fecal, endoscopic and X-ray examinations | 3 | 0.3% | 0.1–1.0 |
| Joint fecal, endoscopic, crop endoscopy and X-ray examinations | 3 | 0.3% | 0.1–1.0 |
| Aspiration of swelling and microscopic examination | 3 | 0.3% | 0.1–1.0 |
| Crop endoscopy and X-ray | 2 | 0.2% | 0.0–0.8 |
| Fecal examination and X-ray | 2 | 0.2% | 0.0–0.8 |
| Joint endoscopic and fluorescent eye stain tests | 2 | 0.2% | 0.0–0.8 |
| Joint endoscopic and biochemistry tests | 2 | 0.2% | 0.0–0.8 |
| **Total** | **906** | **100%** |  |

### Incidence of the different health problems of falcons treated at City Vet Clinic

The overall incidence of the different health problems of falcons identified by the City Vet clinic in 906 falcons visited the clinic during 2019 was 26.5% (95% confidence interval, CI, 23.6, 29.5%). Ingluvitis, inflammation of the crop, was the most frequently occurring health problem in falcons visited City Vet Clinic in 2019 with incidence rate of 8.1% (95% CI, 6.4, 10.0%). Next to Ingluvitis, aspergillosis was the second most frequent incident disease that affected Falcons treated at City Vet Clinic in 2019 and the incidence of aspergillosis was 5.8% (95% CI, 4.4, 7.6%) (Table 3). Bacterial enteritis was the 3^rd^ most incident health problem of Falcons visiting the City Vet Clinic in 2019 and its incidence was 2.4% (95% CI, 1.5, 3.7%).

**Table 3.** Incidence of the major health problems of 906 falcons diagnosed at the City Vet Clinic, Al Ain City in 2019.

| **Health problem** | **Number affected** | **Incidence of the health problem** | **95% Confidence interval** |
| --- | --- | --- | --- |
| Inguvitis, alone or with co-morbidities | 73 | 8.1% | 6.4–10.0 |
| Aspergillosis alone or with co-morbidities | 53 | 5.8% | 4.4–7.6 |
| Coccidiosis alone or with co-morbidities | 15 | 1.7% | 0.9–2.7 |
| Bacterial enteritis alone or with co-morbidities | 22 | 2.4% | 1.5–3.7 |
| Eye abnormalities (ocular ulcer, inflammation) | 12 | 1.3% | 0.7–2.3 |
| Strigea | 8 | 0.9% | 0.4–1.7 |
| Tapeworm | 6 | 0.7% | 0.2–1.4 |
| Injury to different body parties | 12 | 1.3% | 0.7–2.3 |
| Wound | 9 | 1.0% | 0.5–1.9 |
| Inflammation of the air sac | 7 | 0.8% | 0.3–1.6 |
| Severe dehydration | 6 | 0.7% | 0.2–1.4 |
| Foot problems | 5 | 0.6% | 0.2–1.3 |
| Congestion of lungs | 4 | 0.4% | 0.1–1.1 |
| Trichomoniasis | 4 | 0.4% | 0.1–1.1 |
| Skin lesion | 2 | 0.2% | 0.0–0.8 |
| Fat deposit on the liver | 2 | 0.2% | 0.0–0.8 |
| **Total** | **240** | **26.5%** | **23.6–29.5** |

### Number of falcons visited City Vet Clinic during each Month of 2019

The number of falcons visited City Vet Clinic during the 12 months of 2019 is depicted in Figure 1. The relationship between the number of the cases and the months was not linear; but it was rather polynomial. Quadratic curve was able to fit the data better than the linear line and the value of polynomial regression (R²) was 42% (equation of curve: y = 3.1933x^2^ - 38.541x + 153.05), which implied that only 42% of the variation in number of monthly cases was explained by monthly variations. As it can be seen from Figure 1, the number of cases visited the Clinic during January was high and thereafter the number of cases started to decrease and reached the lowest level in April and stayed low until August when it started to rise and stayed high from August to December.

**Figure 1.**
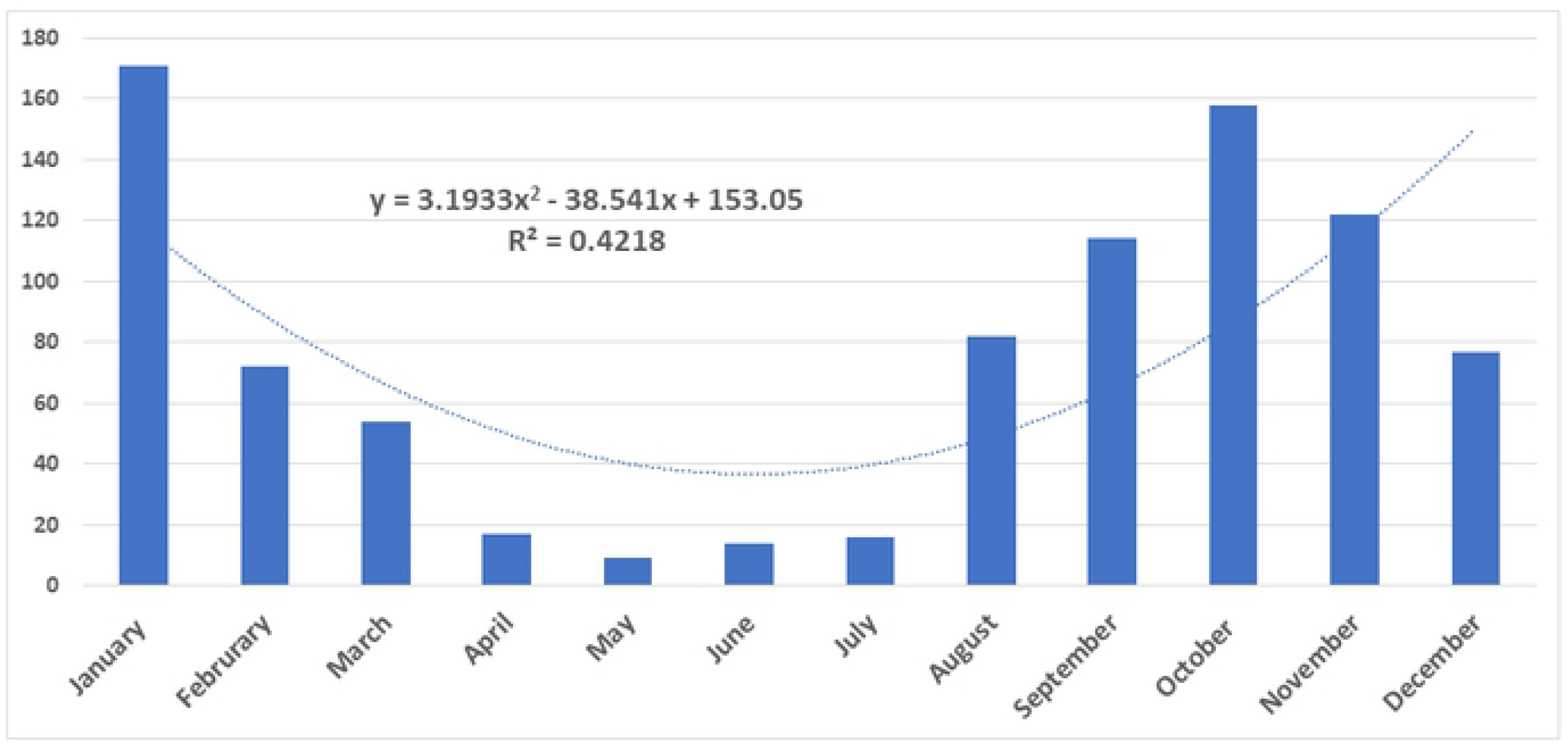
Number of different cases of falcon visited City Vet Clinic in Al Ain in 2019. The relationship between the number of the cases and the months was not linear; but it was rather polynomial and the value of polyno111ial regression (R2) was 42%, which i111plied that only 42% of the variation in nu111ber of cases was explained by 111onths of the Year.

### Types of treatment given to falcons with different health problems

The different types treatment used to treat falcons at the City Vet Clinic are presented in Table 4. Antibiotics were used for the treatment of 46.3% (95% CI: 39.8, 52.8%) of the total cases. Next to antibiotics, antifungal drugs were the second most commonly used drugs and used for the treatment of 21.7% (95% CI: 16.6, 27.4%).

**Table 4.** Types of treatments given to 240 falcons with health by the City Vet Clinic in 2019.

| Type of treatment | Number treated | Percent (%) Treated | 95% Confidence interval |
| --- | --- | --- | --- |
| Antibiotics | 111 | 46.3 | 39.8–52.8 |
| Anti-fungal drugs | 52 | 21.7 | 16.6–27.4 |
| Anthelmintic and anti-coccidian | 29 | 12.0 | 8.2–16.9 |
| Treatment of injuries and wounds | 26 | 10.8 | 7.2–15.4 |
| Eye ointment | 12 | 5.0 | 2.6–8.6 |
| Euthanasia | 6 | 2.5 | 0.9–5.4 |
| Not treated | 4 | 1.7 | 0.5–4.2 |
| **Total** | **240** | **100.0%** |  |

The other types of treatments and services provided to falcons at the City Vet Clinic are presented in Table 5. These include supportive treatments such as rehydration and administration of vitamins. Over 13% of the total falcon were rehydrated while tail mounting and feather fixing were done for 11. 9% (95%CI: 9.9, 14.2%) and 8.7% (95% CI: 7.0, 10.8%), respectively.

## Discussion

The present study was conducted on the health problems of falcons treated at the City Vet Clinic in Al Ain, UAE in 2019. Computerized case record of the Clinic was used for the data extraction for 906 falcons regarding the purpose of visit to the clinic, the diagnosis method used, the health and related problem diagnosed, the type of treatment and other services given to these falcons. The result indicated that the majority of falcons (68.9%) were brought to the Clinic for the checkup of their health conditions. Health checkup is considered as important procedure by the falconers and each falcon is expected to visit a hospital or a clinic twice a year for checkup [6]. In 2004, pre-purchase health checkup has been introduced by the Abu Dhabi Falcon Hospital and as a result, falconers consider health checkup as part of the purchasing procedure [6]. For the purpose of health checkup, combinations of different diagnostic tests were used in order to identify any health problem affecting falcons. The most widely used diagnostic methods were fecal and crop endoscopic examinations. Crop endoscopic examination is restricted to the examination of the crop, which is exposed to infections since it stores feed while fecal examination is used for the diagnosis of infections of the lower gastrointestinal tract. Parasitic infections including protozoan and helminth parasites and bacterial infections were diagnosed by fecal examination. In addition to crop endoscopy, endoscopic examination of the internal organs, X-ray, biochemistry and blood tests were used for the health checkup and diagnosis of diseases of falcons

Following health checkup, tail mounting and feather fixing were the 2^nd^ and 3^rd^ most frequent reasons why falcons were brought to the City Vet Clinic. In tail mounting, a radio transmitter is fixed on the feathers of the tail of a falcon so as to monitor the movement of the falcon. Similarly, feather fixing was also another important reason why falcons visiting City Vet Clinic. Normal feathers are required for falcons to fly well and catch preys. Feathers can easily break when falcons are fighting with the prey or during landing with a high speed and hence require fixing.

From among disease conditions mainly caused by infections, ingluvitis was the most incident (8.1%) health problem that was detected in the study falcons. It was detected by the use of crop endoscopic examination and the etiologic agents were not identified. Ingluvitis can be caused by different infections and or physical damage including overfeeding (the food would stay in the crop for too long and so pathogens would grow and populate inside), dehydration and hypoglycemia, viral diseases, parasites, mycotic diseases and bacterial diseases [13]. A study conducted in the Abu Dhabi Falcon Hospital earlier indicated that 5% of the crops of falcons were infected with *Candida* species and 1.4% were infected by *Trichomonas* [14]. The result of a similar study conducted on the clinical records of 3376 falcons in the Kingdom of Saudi Arabia indicated infectious diseases, traumatic injuries, toxicosis, and metabolic or nutritional diseases are the major health problems of falcons [15].

Next to ingluvitis, the second most incident (5.8%) health problem identified in falcons treated at the City Vet Clinic was aspergillosis or and its co-morbidities. Similar to the present observation, a study conducted in Dubai reported pathological findings associated with *Aspergillus* species in 94 clinically diseased captive falcons [16]. Thus, aspergillosis is considered the most common systemic mycosis in birds and can lead to death in captive falcons [17,18,19]. Aspergillosis affects the lower respiratory tract and isolated from the air sacs of falcons [20]. *Aspergillus* species are widespread in the environment and become pathogenic mainly under stressful conditions such as poor ventilation, malnutrition, toxins, vaccinations, long-term use of antibiotics and corticosteroids, hot-humid climate, and stress-associated conditions, such as recent capture, training, and change of ownership [21,22,23].

Bacterial enteritis was recorded as the 3^rd^ most frequent health problem of falcons visited the City Vet Clinic in Al Ain in 2019. Bacterial enteritis is usually caused by *Escherichia coli* (*E. coli*) and *Salmonella* species. Earlier study indicated that among bacterial isolates, *E. coli* was isolated from 35% of 663 diseased raptors with clinical signs of septicemia or respiratory disorders while *Pseudomonas aeruginosa* was isolated from 7% of the same group of raptors [24]. Another study indicated that the common bacterial diseases in falcons are chlamydiosis, salmonellosis, avian tuberculosis and mycoplasmosis [25]. However, in the present study, bacterial identification was not performed and hence, identification of the bacteria using standard procedures is required in order to detect the bacterial species affecting falcons in Al Ain.

The incidence of infection with Coccidia was about 2% and it was in the order of the important infections. Similar to the observation of the present study, coccidiosis was reported in raptors earlier [26]. From among the coccidian parasites, the Genus *Caryospora* infects predatory birds and reptiles [27]. According to the existing literature, least 25 species of *Caryospora* have been identified from birds worldwide [28] of which 15 have been identified from raptors [29–31]. However, in the present study, species level identification has not been made and hence a better designed and more detail study is required on coccidiosis in falcons in Al Ain and UAE.

Viral diseases have not been recorded on the case records of the Clinic. However, vaccination was administered to about 10% of the falcons visited the Clinic in 2019 against avian flu, Newcastle disease and avian pox. These three diseases are important viral diseases of birds and have also been reported in falcons. For example, Newcastle disease has been reported to be an important viral disease of falcons in the Middle East [32]. Besides, Avian flu, particularly the highly pathogenic avian influenza A (HPAI) virus is a major threat to the avian species and humans in the worldwide [33]; and in 2000, H7N3 was isolated from a Peregrine falcon (*Falco peregrinus*) kept as a falconry bird in UAE [34]. Additionally, during the outbreak of H7N7 in poultry in Italy, H7 subtype was isolated from a Saker falcon (*Falco cherrug*) [35]. Similarly, experimental infection of falcons with H5N1 lead to death of infected falcons [36]. The isolation of HPAI from falcons has a very serious public health implication as falconers have close physical contacts with their falcons. Hence, regular vaccination of falcons against Avian Influenza A should be considered priority. Similar to Avian flu and Newcastle disease, Poxviruses were also reported in species of falconidae imported to Germany from Arabian or Asian countries [12].

Helminth parasites such as *Strigea* and *Tapeworm* were detected in the feces of falcons with the incidence rates of about 1% each. These parasites can cause different problems such as anemia, weight loss, loss of appetite, diarrhea, and even death in severe cases. Therefore, regular checkup and deworming are required in order to control parasitic infections in falcons.

## Conclusion

The result of this study showed that majority of falcons visited the City Vet Clinic for health checkup following which tail mounting and feather fixing were the second and third purposes for visit to the Clinic, respectively. The identified major health problems of falcons were ingluvitis aspergillosis, bacterial enteritis, coccidiosis, injuries and eye abnormalities in decreasing frequency of occurrence. These health problems can threaten falcons and can also be transmitted to owners. Therefore, regular checkup of falcons, and control of diseases of falcons were recommended.

## Abbreviations

HPAI: Highly pathogenic avian influenza A virus
H7N3: Hemagglutinin 7 and Neuraminidase 3
H5N1: Hemagglutinin 5 and Neuraminidase 1
UAE: United Arab Emirates

## Acknowledgments

The owner of the City Vet Clinic (Al Ain) is highly appreciated for giving us access to the clinic data and collaborating with us working on the data. The other staff of the Clinic are acknowledged for the support during data collection.

## Authors’ contributions

MAH performed data extraction and drafted the manuscript. AAd supported in data extraction and edited the manuscript. KM assisted in designing the study and edited the manuscript. BAD edited the manuscript. GA designed the study, analyzed the data and edited the manuscript.

## Ethical approval and consent to participate

The study was approved by the Council of the Department of Veterinary Medicine of the College of Food and Agriculture of the UAE University as a Senior student research Project of the first author. Additionally, the study obtained support from the City Vet Clinic.

## References

1. Cenizo, M., Noriega, J.I., Reguero, M.A. A stem falconid bird from the Lower Eocene of Antarctica and the early southern radiation of the falcons. J. Ornithol. 2016; 157: 885–894

2. Gill, F. and Brown, L. H. 2021. Falconiform. In: Encyclopaedia Britanica, Inc. [Internet] [cited on 8 February 2021]. Available: https://www.britannica.com/animal/falconiform

3. Rodriguez, E., Deubel, J., Gaur, A., Singh, S., Promeet, D. 2021. Falcon, Bird. In: Encyclopaedia Britannica. [Internet] [cited on 8 February 2021]. Available: https://www.britannica.com/animal/falcon-bird

4. Sultan Al-Ulama, Mohammad Ismail M. A. 1997. Study on the Parasites of Falcons in the United Arab Emirates. MSc Theses 665. Available: https://scholarworks.uaeu.ac.ae/all_theses/665

5. Wallen K.E. and Bickford N.A. Stakeholder perspectives on raptor conservation and falconry in North America. Glob. Ecol. Conserv. 2020; 24: e01280

6. Muller, M.G. 2009. Practical Handbook of Falcon Husbandry and Medicine. Nova Publishers Inc. New York, ISBN: 978-1-60741-608-1 Available upon request

7. Muller, M.G., Mannil, A.T. and A. George. 2009. Study on the most common bacterial infections in falcons in the United Arab Emirates. Abu Dhabi Falcon Hospital/Environmental Research and Wildlife Development Agency (ERWDA), P.O. Box 45553, Abu Dhabi, United Arab Emirates https://www.falconhospital.com/media/1459/common-bacteria-1.pdf

8. Townsend, S. 205. “Sheikh Hamdan’s Bid to Revive the Glorious Arab Sport of Falconry.” In: Arabian Business. [Internet] [cited on February 8, 2021]. Available: https://www.arabianbusiness.com/sheikh-hamdan-s-bid-revive-glorious-arab-sport-of-falconry-595769.html

9. Lotha, G. 2019. Bedouin. In: Encyclopaedia Britannica, Inc. [Internet] [cited on 8 February 2021]. Available: https://www.britannica.com/animal/falconiform

10. Jacobs, H. 2020. Middle-east-falcons-uae-training-2019-1. In: Business Insider. [Internet] [cited on 12 July 2020]. Available: https://www.businessinsider.com

11. Mass Audubon, 2021. Common Bird Parasites & Diseases. [Internet] [cited on 8 February 2021]. Available: https://www.massaudubon.org/learn/nature-wildlife/birds/common-bird-parasites-diseases

12. Krone, O., Essbauer, S., Wibbelt, G., Isa, G., Rudolph, M., R. E. Gough, R.E. Avipoxvirus infection in peregrine falcons (Falco peregrinus) from a reintroduction program in Germany. Vet. Rec. 2004; 154: 113–114

13. Zucca, P. and Delogu, M. 2008. Infectious diseases. In: Samour, J. (ed). Avian Medicine. 2nd ed. Mosby Elsevier. pp. 309–337.

14. Muller, M.G. and Nafeez, M.J. 2004. Pre-purchase examinations in first year captivebred falcons; Wildlife Diseases Association Conference, 11th - 13th December 2004, Abu Dhabi. Compiled on CD Rom by Mwanzia, J. and P. Soorae.

15. Naldo, J.L. and Samour, J.H. Causes of Morbidity and Mortality in Falcons in Saudi Arabia. J. Avian Med. Surg. 2004; 18(4): 229–241

16. Tarello, W. Etiologic Agents and Diseases Found Associated with Clinical Aspergillosis in Falcons. Int. J. Microbiol. 2011, 6 pages, doi:10.1155/2011/176963

17. Cooper, J.E. 1985. Veterinary Aspects of Captive Birds of Prey, Standfast Press, Cherington, UK, 2nd edition, 1985.

18. Kunkle, R. A. and Rimler, R.B. Early pulmonary lesions in turkeys produced by nonviable Aspergillus fumigatus and/or Pasteurella multocida lipopolysaccharide. Avian Dis. 1998; 42(4): 770–780

19. Samour, J. H. 2000. Veterinary considerations during the hunting trip. In: Raptor Biomedicine III, J. T. Lumeij, J. D. Remple, P. T. Redig, M. Lierz, and J. E. Cooper, Eds., pp. 267–274, Zoological Education Network, Lake Worth, Fla, USA.

20. Silvanose, C.D., Bailey, T.A., and Di Somma, A. Susceptibility of fungi isolated from the respiratory tract of falcons to amphotericin B, itraconazole and voriconazole. Vet. Rec. 2006; 159(9): 282–284.

21. Deem, S. L. Fungal diseases of birds of prey. Vet. Clin. North Am. Exot Anim. Pract. 2003; 6(2): 363–376

22. Redig, P. 2003. Aspergillosis. In: Avian Medicine, J. Samour, Ed., pp. 275–287, Elsevier Science Limited, Edinburgh, Scotland

23. Mukaratirwa, S. Outbreak of disseminated zygomycosis and concomitant pulmonary aspergillosis in breeder layer cockerels. J. Vet. Med. B. 2006; 53(1): 51–53

24. Vidal, A., Baldomà, L., Molina-López, R.A., Martina, M., and Darwich, L. Microbiological diagnosis and antimicrobial sensitivity profiles in diseased free-living raptors. Avian Pathol. 2017; 46(4): 442–450 https://doi.org/10.1080/03079457.2017.1304529

25. Zubair, M. 2004. Biology and Behavior of Falcons with Emphasis on Captive Breeding and Healthcare of Peregrine Falcon (Falco peregrinus). MSc Thesis. Department of Zoology, Farook College Calicut, University of Calicut, Kerala.

26. Klaphake, E. and Clancy, J. 2005. Raptor Gastroenterology. Vet. Clin. North Am. Exot. Anim. Pract. 2005; 8: 307–327.

27. Upton, S.J., Current, W.L., Barnard, S.M., 1986. A review of the genus Caryospora Leger, 1904 (Apicomplexa: Eimeriidae). Syst. Parasitol. 1986; 8: 3–21.

28. Yang, R., Brice, B. and Una Ryan, U. 2014. A new *Caryospora* coccidian species (Apicomplexa: Eimeriidae) from the laughing kookaburra (*Dacelo novaeguineae*). Exp. Parasitol. 2014; 145: 68–73

29. Upton, S.J., Campbell, T.W., Weigel, M., McKown, R.D. The Eimeriidae (Apicomplexa) of raptors: review of the literature and description of new species of the genera Caryospora and Eimeria. Can. J. Zool. 1990; 68: 1256–1265.

30. Alfaleh, F.A., Alyousif, M.S., Al-Shawa, Y.R., Al-Quraishy, S. Caryospora cherrughi sp. (Apicomplexa: Eimeriidae) infecting *Falco cherrug* in Saudi Arabia. Parasitol. Res. 2013; 112: 971–974.

31. McAllister, C.T., Duszynski, D.W., McKown, R.D. A new species of Caryospora (Apicomplexa: Eimeriidae) from the sharp-shinned hawk, Accipiter striatus (Aves: Accipitriformes). J. Parasitol. 2013; 99: 490–492.

32. Samour, J. 2014. Newcastle Disease in Captive Falcons in the Middle East: A Review of Clinical and Pathologic Findings. J. Avian Med. Surg. 2014; 28(1): 1–5, doi: 10.1647/2011-041.

33. Lierz, M., Hafez, H.M., Klopfleisch, R., Lüschow, D., Prusas, C., Teifke, J. P., et al. Protection and Virus Shedding of Falcons Vaccinated against Highly Pathogenic Avian Influenza A Virus (H5N1). Emerg. Infec. Dis. 2007; 13 (11): 1667–1674

34. Manvell RJ, McKinney P, Wernery U, Frost K. Isolation of a highly pathogenic influenza A virus of subtype H7N3 from a peregrine falcon. Avian Pathol. 2000; 29: 635–7.

35. Magnino S, Fabbi M, Moreno A, Sala G, Lavazza A, Ghelfi E, et al. 2000. Avian influenza virus (H7 serotype) in a saker falcon in Italy. Vet. Rec. 2000;146: 740.

36. Bertran K, Busquets N, Abad FX, Garc’a de la Fuente J, Solanes D, et al. 2012. Highly (H5N1) and Low (H7N2) Pathogenic Avian Influenza Virus Infection in Falcons Via Nasochoanal Route and Ingestion of Experimentally Infected Prey. PLoS One. 2012; 7(3), e32107. doi:10.1371/journal.

